# Neural basis of perceptual surface qualities: Evidence from EEG decoding

**DOI:** 10.1101/2024.02.05.578885

**Authors:** Taiki Orima, Suguru Wakita, Isamu Motoyoshi

## Abstract

The human visual system can easily recognize object material categories and estimate surface properties such as glossiness and smoothness. A number of psychophysical and computational studies suggest that the material perception depends on global feature statistics of the entire image at multiple processing levels. Neural representations of such global features, which is independent of precise retinotopy, may be captured even by EEG that have low spatial resolution. To test this possibility, here we measured visual evoked potentials (VEPs) for 191 natural images consisting of 20 categories of materials. We then sought to classify material categories and surface properties from the VEPs, and to reconstruct the rich phenomenological appearance of materials themselves via neural representations of global features as estimated from the VEPs. As a result, we found that material categories were correctly classified by the VEPs even at latencies of 150 ms or less. The apparent surface properties were also significantly classified within 175 ms (lightness, colorfulness, and smoothness) and after 200 ms (glossiness, hardness, and heaviness). In a subsequent reverse-correlation analysis, we further found that the VEPs at these latencies are highly correlated with low- and high-level global feature statistics of the surface images; Portilla-Simoncelli texture statistics and style information in deep convolutional neural network (dCNN), indicating that neural activities about such global features are reflected in the VEPs that enabled successful classification of materials. To demonstrate this idea more directly, we trained deep generative models (MVAE models) that reconstruct the surface image itself from the VEPs via style information (gram matrix of the dCNN output). The model successfully reconstructed realistic surface images, a part of which were nearly indistinguishable from the original images. These findings suggest that the neural representation of statistical image features, which were formed at short latencies in the visual cortex and reflected even in EEG signals, not simply enable human visual system to recognize material categories and evaluate surface properties but provides the essential basis for rich and complex phenomenological qualities of natural surfaces.

## Introduction

Objects in the natural environment have unique surface properties derived from a wide variety of materials, and the retinal images of these natural surfaces are extremely complex. However, we can easily estimate surface properties such as glossiness, transparency, and hardness (Motoyoshi et al., 2007; Sharan et al., 2008; Fleming et al. Kentridge, 2015; Fleming, 2017) and can rapidly and reliably recognize material categories (Fleming et al., 2013; Wiebel et al., 2013; Sharan et al., 2013, 2014; Fleming, 2017). A large number of psychophysical studies have revealed how the visual system correctly or incorrectly estimates material and surface properties from retinal images based on low- and high-level image features.

A series of electrophysiological and fMRI studies have also unveiled neural substrates of visual material and surface property processing (Edward et al., 2003; Cant et al. Hiramatsu et al., 2011; Nishio et al. 2012; Okazawa et al. 2012; Wiebel et al. 2014). For example, Hiramatsu et al. (2011) used fMRI to measure brain activity for CGs mimicking various materials and showed that information related to material perception is processed in a hierarchical manner. Specifically, the dissimilarity matrices of brain activity in the early visual cortex (V1 and V2) strongly correlated with that of image statistics, while the dissimilarity matrix of spindle gyrus activity strongly correlated with that of ratings for each stimulus. These results suggested that the low-level visual cortex represented information about image statistics, while the higher-level visual cortex represented some higher-level information that highly correlated with the perceived surface properties. Another fMRI study on macaque monkeys using similar visual stimuli also suggested that V1 reflects the encoding of low-level image features, while V4 integrates information directly related to the perception of object material properties (Goda et al., 2014). These studies suggest that hierarchical neural representations of low- and high-level image features in the visual cortex support material discrimination and evaluation of surface properties.

In most everyday situations, material perception largely (but not entirely) depends on the image appearance of interior textural regions rather than on the contours of objects. A large body of evidence suggest that the appearance of these textural regions is based on global statistical features of the image region of interest (Bergen & Adelson, 1988; Heeger & Bergen, 1995; Zipser et al. Portilla & Simoncelli, 2000; Baker & Mareschal, 2001; Motoyoshi et al. 2007; Freeman & Simoncelli, 2011; Freeman et al. 2013; Ziemba et al. 2019; Orima & Motoyoshi, 2021; Okada & Motoyoshi, 2021). For example, the best-known model by Portilla & Simoncelli (2000) can predict the appearance of various natural texture images by energy, autocorrelation/cross-correlation, and moment statistics of subband images of different orientations and spatial frequencies, and eventually can synthesize perceptually equivalent texture images. Recent neurophysiological studies have shown that these low-level statistical features represented in the PS models are encoded in V1 and V2 (Freeman & Simoncelli, 2011; Freeman et al. 2013; Okazawa et al. 2015, 2017; Ziemba et al. 2019), and the higher-level statistical features constituted by the integration of low-level features are represented in V4 (Arcizet et al., 2008; Okazawa et al., 2015, 2017). Furthermore, the neural style transfer (NST) technique (Gatys et al., 2015) focusing on the gram matrix (covariance between channels) in deep convolutional neural networks (dCNN), although less formulated than the previous texture models such as the PS model, describes the appearance of natural texture images much more precisely than the PS model based on statistical feature representations in dCNNs, and the NST can synthesize an almost indistinguishable images from the original images even when they are observed foveally. These findings suggest that the appearance of texture images which is captured at a glance can almost completely be explained by the hierarchical neural representation of low- and high-level statistical image features in the visual cortex.

It has been suggested that human perception of surface materials can be explained by both low- and high-level statistical features of surface areas. For example, some early studies suggested that surface gloss perception is supported by low-level statistics such as moment statistics (Adelson, 2001; Motoyoshi et al. 2007; Sharan et al. 2008; Motoyoshi & Matoba, 2012; Wiebel et al., 2015). However, this idea had obvious limitations (Anderson & Kim, 2009; Wiebel et al., 2015), which was represented by the fact that the PS model failed to reproduce gloss perception perfectly (Wang et al., 2013). Other studies have denied the use of image statistics and instead indicated the importance of the visual cues corresponding to physical properties such as local 3D shapes and highlights (Anderson & Kim, 2009; Kim & Anderson, 2010; Marlow & Anderson, 2013). However, these features are subjectively defined by researchers rather than objectively measured from images, and their experimental evidence were thought to be obtained by using visual stimuli which appear rarely in our daily lives. On the other hand, recent findings, such as the remarkable success of the NST (Gatys et al., 2015) and the recent accurate prediction of gloss perception and misperception using unsupervised deep learning models (Storrs et al., 2021), strongly support the idea that low- and high-level statistical representations for images can still explain the perception of a wide range of material properties. High-level neural activity that could not be explained by image statistics in the previous fMRI studies (e.g., Goda et al., 2014) may be explained by these higher-level statistical features.

Given that the statistical representation is invariant to the spatial layout of features within an image region, we expected that the neural response of these features could be extracted even from EEG with low spatial resolution. In fact, we have shown not only that the visual evoked potentials (VEPs) for various natural texture images were systematically correlated with low-level image statistics, but that we could synthesize perceptually similar textures based on image statistics reconstructed from the VEPs (Orima & Motoyoshi, 2021). In the subsequent study, we also succeeded in synthesizing photorealistic texture images from VEPs using a deep generative model (MVAE) (Wakita et al., 2021). Based on these findings, by using EEG signals, it is natural for us to assume that we can decode material perception, which is believed to be governed by low- and high-level statistical features. If it is possible to decode perceived material categories and surface properties from EEG signals that are evoked while the observation of visual stimuli for a few hundred milliseconds, and to synthesize similar image appearances from the EEG signals, it would further support the idea that a large part of material perception depends on low- and high-level textural (statistical) information.

In the present study, we conducted a series of experiments and analyses using EEG to find dynamic and multidimensional neural representations that support the perception and discrimination of natural surfaces in a single glance. In Part 1, we trained a support vector machine (SVM) using measured VEPs as input. As a result, material categories and perceived surface properties were classified at a statistically significant level, and the temporal development of visual material processing was revealed. These results suggest that neural representations, as reflected in the temporal patterns of the EEG, support the discrimination of natural surface materials and the perception of their characteristics. In Part 2, to clarify how such neural representations are related to statistical features in surface images, we applied representation similarity analysis between the VEPs and subband image statistics or style information in dCNN models. We found that the style information is highly correlated with the VEPs. Inspired by the strong correlation, in Part 3, we attempted to reconstruct the surface images from the VEPs using multimodal variational autoencoder (MVAE) models, which improved on the model in our previous study (Wakita et al., 2021). The results showed that some of the reconstructed images strikingly reproduced the impression of the original images and most of the reconstructed images from the VEPs were similar to the original images. Psychophysical experiments showed that more than 80% of the reconstructed images were more similar to the original than randomly reconstructed images, and that the perceived material categories and perceived surface properties of the reconstructed images matched those of the original images at a statistically significant level. These results support the findings of a series of material perception studies and suggest that even rich and complex properties of object surfaces that are difficult to describe can be decoded by simple VEPs. Therefore, these results further support the idea that human texture and material perception relies on statistical, global features that are reflected even in VEPs.

## Part 1: Classification of Material Categories and Surface Properties from VEPs

First, we measured VEPs for a wide variety of natural surface images made of various materials, and used SVM to examine how the VEPs reflect neural information that supports perceptual categorization of material categories and surface properties.

### Materials & Methods

#### Observers

Fifteen naïve students participated in the experiment. The number of observers was determined according to our previous study (Orima & Motoyoshi, 2021). All observers had normal or corrected vision. All experiments were conducted in accordance with the guidelines set by the “Ethical Review Committee for Experimental Research on Human Subjects, Faculty of Arts and Sciences, The University of Tokyo,” and all observers provided written informed consent.

#### Apparatus

In EEG recordings, the visual stimuli were generated by a PC (HP Z2 Mini G4 Workstation) and presented on a 24-inch gamma-corrected LCD (BenQ XL2420T) with a refresh rate of 60 Hz and a spatial resolution of 1.52 min/pixel at the 88 cm viewing distance we used.

In psychophysical experiments, visual stimuli were generated by a PC (HP Z2 Mini G4 Workstation) and presented on a 24-inch gamma-corrected LCD (BenQ XL2720) with a refresh rate of 60 Hz and a spatial resolution of 1.47 min/pixel at the 73 cm viewing distance we used.

#### Visual stimuli

The original images of the visual stimuli were 191 natural texture images of 256 x 256 pixels (Fig. 1). All images were collected from the Motoyoshi Lab texture database and the Internet. All images were categorized into one of the following 20 material categories: bark, fabric, feather, feather, fish scale, fur, glass, grass, gravel, hair, leather, metal, paper, paste, plastic, powder, rock, skin, soil, sponge and wood. The number of images, the sample size of the following analysis, was determined in accordance with two criteria: more than 8 images in each material category and more than 166 images (Orima & motoyoshi, 2021) in total to retain certain power in statistical tests. These numbers were fixed at the beginning of the present study and never be changed through the whole analysis.

**Figure 1.**
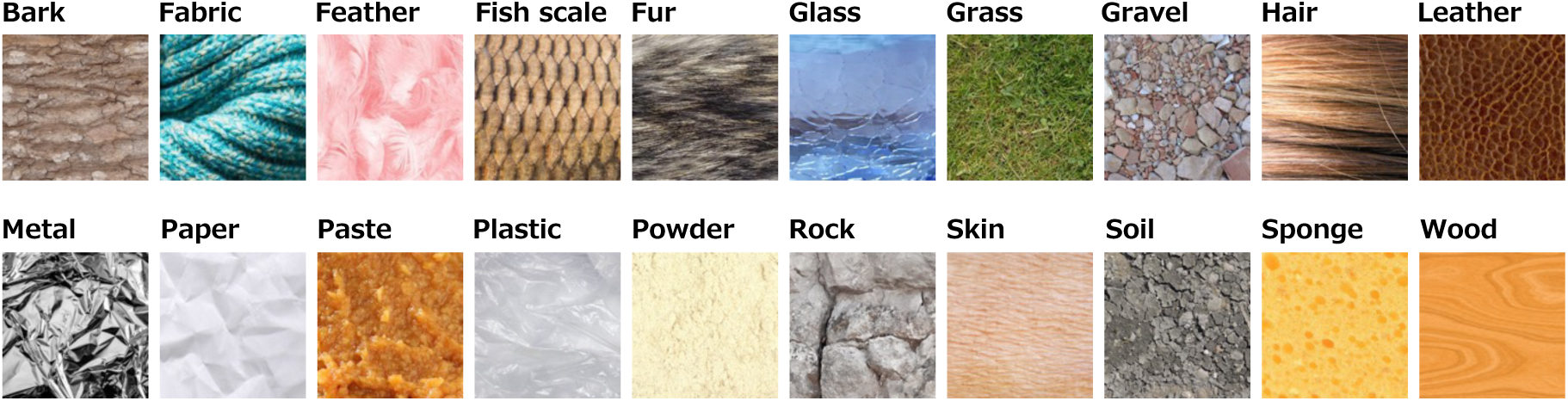
Visual stimuli used in the experiment. One image from each material category is shown as a representative.

Each image was further divided into five variants, each of which was cropped from the upper left, upper right, lower left, lower right, and center of each image to a size of 5.7 x 5.7 deg (224 x 224 pixels), which were used for EEG recordings.

#### EEG recording procedure

The EEG experiments were conducted in a shielded dark room. In each session, 191 of 5.7 x 5.7 deg natural surface images were presented once in a random order. All images were presented for 500 ms, followed by a 750 ms uniform background blank. Observers viewed the visual stimuli foveally through steady fixation on the small black dot that appeared at the center of the screen. EEG recordings were made while the observers viewed the visual stimuli. The observer’s eye movements were controlled by the pre-experiment instruction (cf. Orima & Motoyoshi, 2021, 2023). This session was repeated 5 times for each of 5 variants, 25 times in total for each observer. The same variant was presented in the same session.

#### EEG data preprocessing

EEG data were obtained from 31 electrodes (Fp1, Fp2, F3, F4, C3, C4, P3, P4, O1, O2, F7, F8, T7, T8, P7, P8, Fz, Cz, Pz, FC1, FC2, CP1, CP2, FC5, FC6, CP5, CP6, FT9, FT10, TP9, and TP10) in accordance with the international 10-20 system at a sampling rate of 1000 Hz (BrainVision Recorder, BrainAmp amplifier, EasyCap; Brain Products GmbH). The impedance of each electrode was kept below 5 kΩ (except Cz, P3, and FT10 for one observer). An additional electrode, located between Fz and AFz, was used as a ground electrode. In addition, all electrodes were referenced to another electrode located between Fz and Cz, and all electrode data were re-referenced offline using the average of all electrodes. The recorded EEG data were filtered by a 0.1-40 Hz bandpass filter and divided into epochs of -0.4-0.8 s from the stimulus onset. Baseline correction was performed using -100-0 ms from stimulus onset as a baseline. The artifact component (i.e., eye movements) was removed by independent component analysis and the epochs including abnormal amplitude (exceeding the range from -75 to 75 μV) were rejected to remove epochs with eye blinks. Additionally, 28 epochs (0.04 % of the total) in which the trigger did not work correctly were excluded from the data.

#### Measurement of surface properties values

In a separate session from the EEG experiment, fifteen observers rated the surface properties of each visual stimulus on a 5-point scale from 0 to 4. Observers rated the lightness (dark-light), colorfulness (achromatic-vividly colored), smoothness (rough-smooth), glossiness (matte-gloss), hardness (soft-hard), and heaviness (light-heavy) of a 6.3 x 6.3 deg visual stimulus presented in random order at the center of the screen. The observers rated twice for each visual stimulus with free viewing. The averaged values within each observer and then across observers were used as the representative values of each surface property.

### Results

#### VEPs

Fig. 2 shows the grand average VEPs for all images every 50 ms from 50 ms to 500 ms after the stimulus onset. Red indicates positive amplitudes and blue indicates negative amplitudes. The VEPs were particularly large in the occipital lobe (O1/O2). A rise in the VEPs at the occipital lobe (O1/O2) was observed about 100 ms after the stimulus onset, and then the VEP amplitude began to decrease about 150 ms and later. Thereafter, the amplitude increased again in the occipital lobe, peaking at about 250 ms, and then decreased in the occipital lobe.

**Figure 2.**
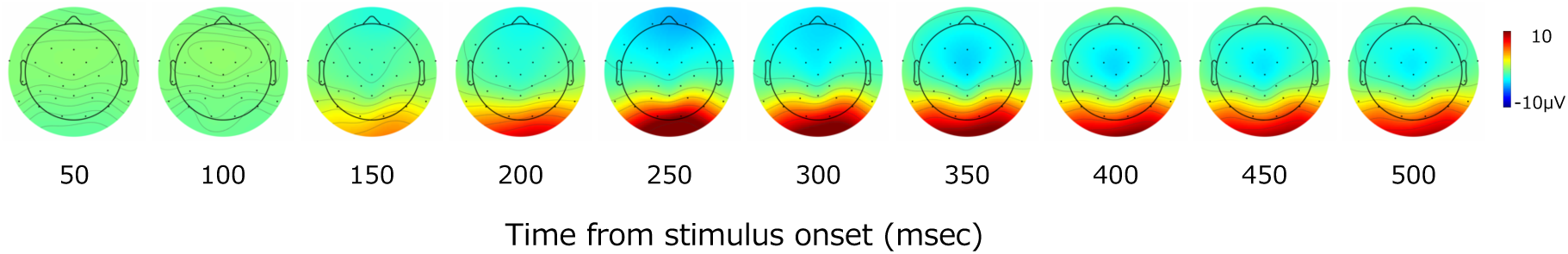
Topographical map of measured VEPs. The color of the map represents the amplitude of the VEPs and corresponds to the indicator on the right.

#### Classification of material category

To examine whether the VEPs reflect the information sufficiently enough to discriminate material categories, we trained SVMs to classify the 20 material categories of the corresponding visual stimuli using the VEPs as input.

EEG data were preprocessed as explained in the method section and 1∼50, 1∼75, …, 1∼500 ms data of the stimulus onset (time-accumulation condition), or 1∼50, 26∼75, …, 451∼500 ms of the stimulus onset were used as input (time-window condition). The EEG data from 29 electrodes (F3, F4, C3, C4, P3, P4, O1, O2, F7, F8, T7, T8, P7, P8, Fz, Cz, Pz, FC1, FC2, CP1, CP2, FC5, FC6, CP5, CP6, FT9, FT10, TP 9, and TP10) was used in the analyses. We used the VEPs averaged across observers as input because the maximum number of VEPs for each visual stimulus for each observer was 25, and even if all of them were averaged, they were still noisy. For each classification, the time and electrode data for each image were flattened, resulting in 29xT (T: number of time points) variables for each image, but the maximum number of independent variables was 14,500, which was much larger than the sample size of 191. Therefore, taking advantage of the fact that the data for each time point and each electrode in the VEPs were correlated with each other, principal component analysis (PCA) was applied to the VEPs, and the principal component (PC) scores up to the PCs whose cumulative explanatory power were greater than 99% were used for the classifications. The number of selected principal components was an average of 123 and a maximum of 144 for the time-accumulation condition, and an average of 58 and a maximum of 73 for the time-window condition, all of which were smaller than the sample size of 191. The classifications were performed using the Matlab function ‘fitcecoc’ as a multiclass classification using the one vs one method.

To perform statistical test of the classifications using the number of images as the sample size, the train/test split in the SVM training was performed using leave-one-out method. In other words, only the VEPs for one image were used as testing data, and the remaining VEPs were input for training the SVM model. The predictions were made by inputting the testing VEP to the trained SVM model. This was repeated 191 times for each SVM model, the number of images, and the average of the prediction was used as the classification accuracy of the SVM model.

Fig. 3 shows the classification accuracy of material categories by the SVM models. The light blue line indicates the classification accuracy of the time-accumulation condition, the yellow line indicates the classification accuracy of the time-window condition, and the error bars indicate the standard error of the classification accuracy at each time. Each colored line at the top of the graph indicates that the corresponding classification accuracy was statistically significant (*p* <= .008) by binomial tests. We adopted the Benjamini-Hochberg (BH) false discovery rate (FDR) correction method (Benjamini & Hochberg, 1995) with α of .05 to address the multiple comparison. The dotted line indicates the chance level (5 %).

**Figure 3.**
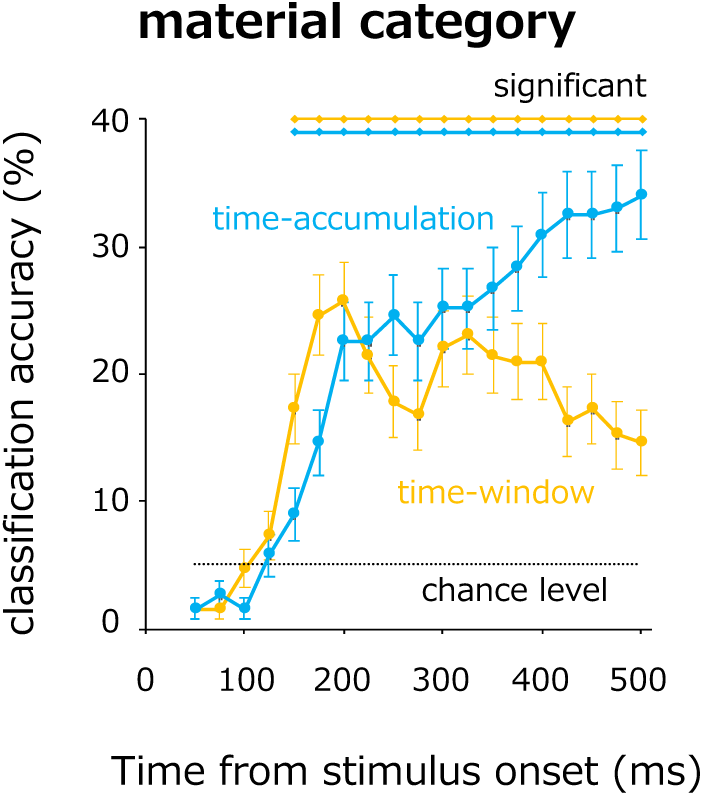
Classification accuracy of 20 material categories by the VEPs. Light blue line represents the classification accuracy of the time-accumulation condition, and yellow line represents the classification accuracy of the time-window condition. Error bars indicate ±1 s.e.m. between cross-validation sets.

Under the time-accumulation condition, the classification accuracy of the 20 material categories became significant at 150 ms after the stimulus onset, eventually reaching nearly 35 %, much higher than the chance level. The results of the time-window condition revealed that the classification accuracy was greatest at around 200 ms after the stimulus onset.

These results suggest that the information encoded before 200 ms of the stimulus onset may contribute significantly to the discrimination of material categories. The classification results of the time-accumulation condition also showed that the classification accuracy continued to increase even at later latencies, such as 300 ms after the stimulus onset, suggesting that information encoded at later latencies also contributed to the encoding of material categories.

#### Classification of perceived surface properties

Next, to examine whether the VEPs reflect the information related to the evaluation of various surface properties, we split rating values of each surface property into 5-steps to distribute the same number of data for each step, and we trained SVMs to classify 5-step degree of each surface property (lightness, colorfulness, smoothness, hardness, and heaviness) of the corresponding visual stimuli using the VEPs as input. The classification was performed using the VEPs up to the certain time point.

EEG data were preprocessed as explained in the method section and 1∼50, 1∼75, …, 1∼500 ms data of the stimulus onset were used as input. The EEG data from 29 electrodes (F3, F4, C3, C4, P3, P4, O1, O2, F7, F8, T7, T8, P7, P8, Fz, Cz, Pz, FC1, FC2, CP1, CP2, FC5, FC6, CP5, CP6, FT9, FT10, TP 9, and TP10) were used in the analyses. We used the VEPs averaged across observers as input because the maximum number of VEPs for each visual stimulus for each observer was 25, and even if all of them were averaged, they were still noisy. For each classification, the time and electrode data for each image were flattened, resulting in 29xT (T: number of time points) variables for each image, but the maximum number of independent variables was 14,500, which was much larger than the sample size of 191. Therefore, taking advantage of the fact that the data for each time point and each electrode in the VEPs were correlated with each other, principal component analysis (PCA) was applied to the VEPs, and the principal component (PC) scores up to the PCs whose cumulative explanatory power were greater than 99% were used for the classifications. The number of principal components selected was an average of 123 and a maximum of 144, all of which were smaller than the sample size of 191. The classifications were performed using the Matlab function ‘fitcecoc’ as a multiclass classification using the one vs one method.

To perform statistical test of the classifications using the number of images as the sample size, the train/test split in SVM training was performed using the leave-one-out method. In other words, only the VEPs for one image were used as testing data, and the remaining VEPs were input for training the SVM model. The predictions were made by inputting the testing VEP to the trained SVM model. This was repeated 191 times for each SVM model, the number of images, and the average of the prediction was used as the classification accuracy of the SVM model.

Fig. 4 shows the classification accuracy of the SVM models classifying surface properties. In each panel, the color of each line corresponds to the legend, with darker colors corresponding to larger rating values. Each colored line at the top of each panel indicates that the corresponding classification accuracy was statistically significant (*p* <= .019, .029, .013, .005, .023, and .013 for lightness, colorfulness, smoothness, glossiness, hardness, and heaviness, respectively) by binomial tests, and the dotted line represents the chance level (20 %). We adopted the BH FDR correction method with α of .05 to address the multiple comparison.

**Figure 4.**
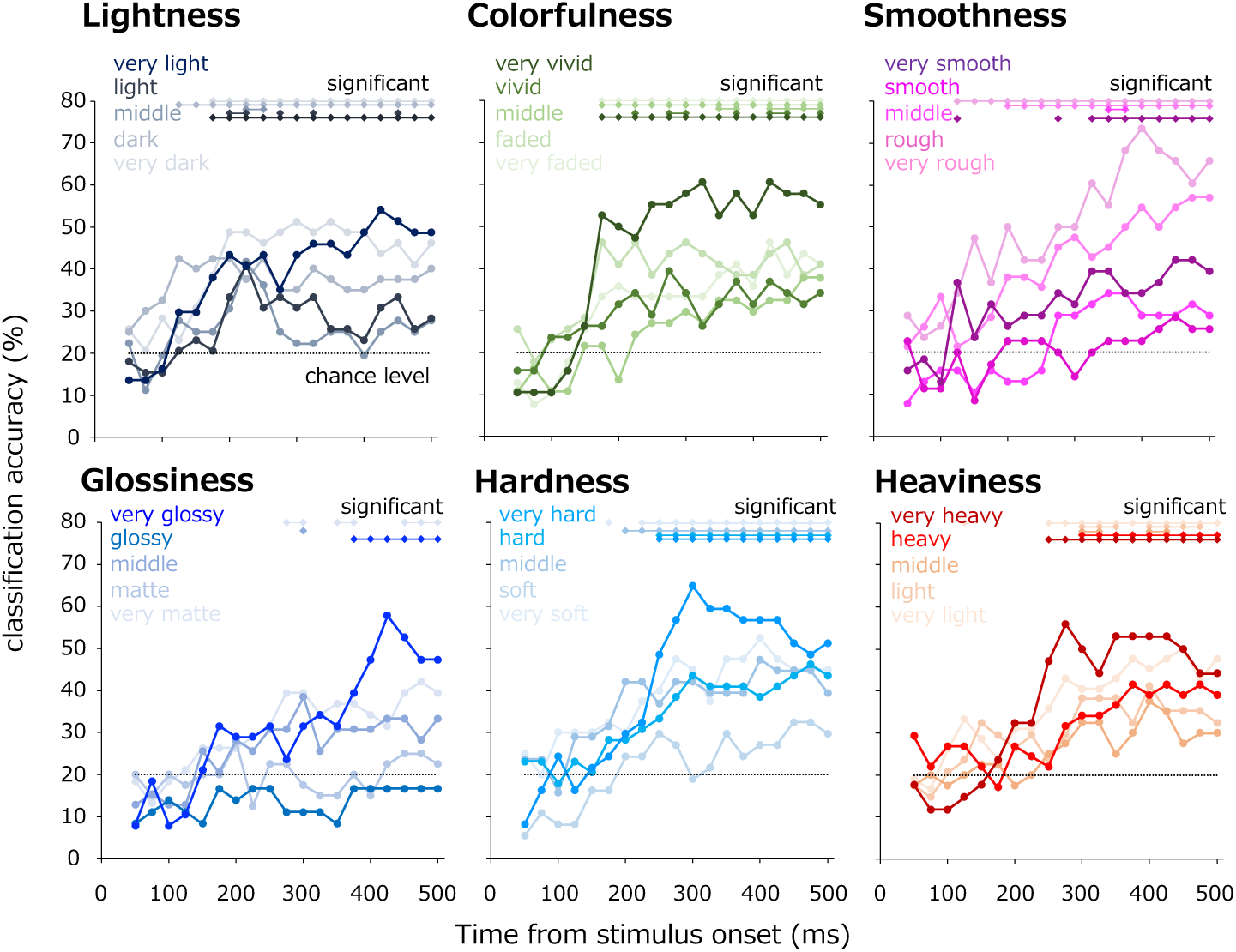
Classification accuracy of surface properties by the VEPs. Each graph represents the classification accuracy when all the VEPs up to the time of each point were input. The error bars corresponding to ±1 s.e.m. between cross-validation sets are omitted for the sake of graph readability, since they are determined dependently from the number of data and the average value.

The classification accuracies of lightness, colorfulness, and smoothness were statistically significant within 175 ms after the stimulus onset, while those of glossiness, hardness, and heaviness became consecutively significant only after 200 ms from the stimulus onset. In addition, the classification accuracies were varied among the surface properties. For instance, smoothness was classified with high accuracy, including the group that was classified with more than 70 %, while glossiness was classified with high accuracy only in the ‘very glossy’ group.

These results suggest that perceived material category, lightness, colorfulness, and smoothness of natural surfaces can be classified by the global information that is captured in the VEPs within 200 ms from the stimulus onset. On the other hand, glossiness, hardness, and heaviness cannot be distinguished by the early latency VEPs.

## Part 2: Relationship between Statistical Image Features and VEPs

The results of Part 1 showed that VEPs for various surface images can classify the perceived material categories and surface properties of the corresponding visual stimuli. Given that VEPs tend to reflect global properties of visual stimuli, these results suggest that the perception of natural surface materials and characteristics are supported by global statistical features. To verify this idea more directly, we investigated how the VEPs for natural surface images correlated with the two representative statistical image features: image statistics that have been suggested to contribute to texture perception in previous studies (e.g., Portilla & Simoncelli, 2000) and style information computed by the neural style transfer technique (Gatys et al., 2015), which is capable to capture rich and complex surface properties.

### Materials & Methods

#### Computation of the image statistics

First, we computed low-level statistical features, image statistics, that have been shown to be important for texture perception in the previous studies (Portilla & Simoncelli, 2000; Hiramatsu et al., 2011; Orima & Motoyoshi, 2021). Each image was converted to Lab color space. For the L channel, the image was decomposed to subband images of 7 spatial frequencies (2- 128 c/image, 1-octave steps) and 8 orientation bands (0-157.5°, 22.5° steps) using bandpass filters of 1-octave spatial frequency bandwidth and 30° orientation bandwidth. For each subband images, the moment statistics (log SD, skewness, and log kurtosis) were calculated. In addition, correlations between subband energy images of different orientation and spatial frequency were calculated. These image statistics were properly summarized in the same manner as our previous study (Orima & Motoyoshi, 2021) considering the nature of textures. For the color channels, pixel moment statistics were calculated according to the previous studies (cf. Hiramatsu et al., 2011). These procedures yielded 47 values of image statistics for each visual stimulus.

The middle row of Fig. 5 illustrates examples of synthesized images generated using the algorithm provided by Portilla & Simoncelli (2000), which synthesizes images using image statistics that are theoretically equivalent to those computed in the present study. The PS synthesized images retain some aspects of surface materials and characteristics, but other surface properties, such as glossiness and hardness, are different from the original image (see Wang et al., 2013).

**Figure 5.**
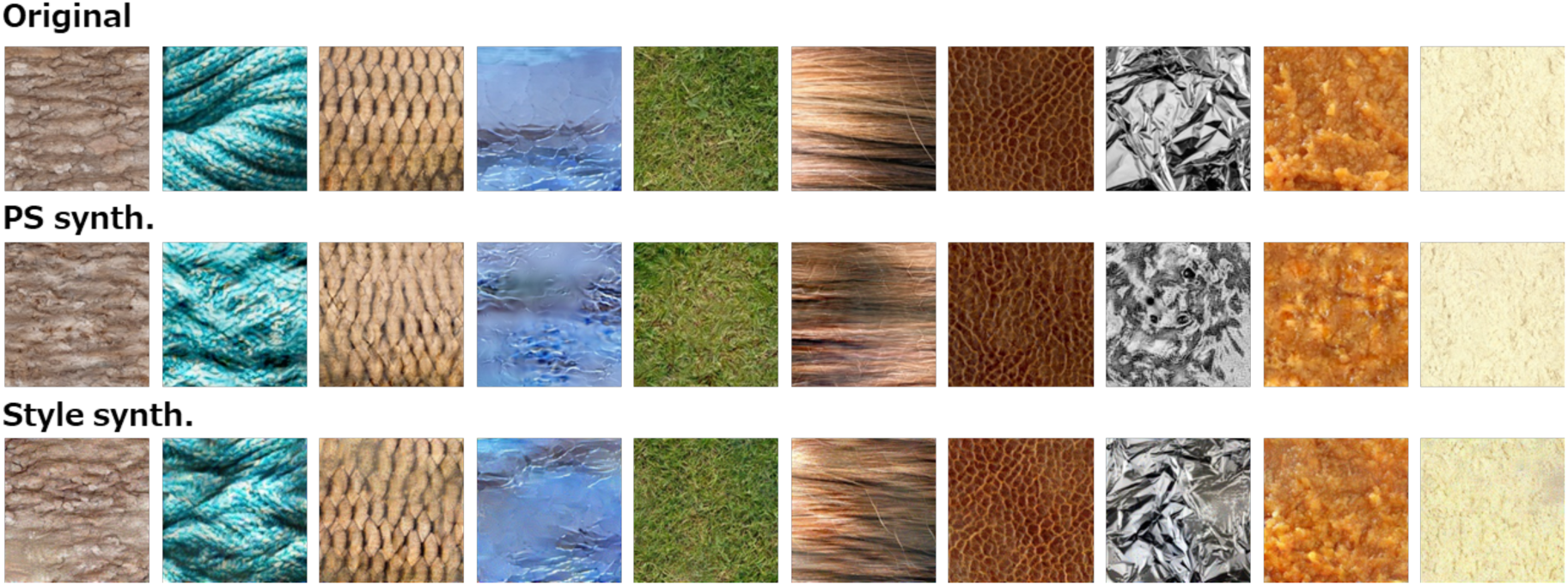
Examples of PS-synthesized and style synthesized images. Each image in the lower row corresponds to the original image in the first row in the same column.

#### Computation of the style information

Neural style transfer (NST) developed by Gatys et al. (2015) is a technique that enables us to apply a “style” of images, which describes complex surface properties such as painting styles of artistic images, to arbitrary images. The style information computed by this algorithm is calculated as a gram matrix of activations of convolutional blocks in a dCNN such as VGG19 (Simonyan & Zisserman, 2014), which performs object recognition with a high accuracy.

In the present study, we computed the gram matrices of the activations of the first to fifth convolutional blocks when each visual stimulus was input to VGG19. The sizes of the gram matrices derived from the first to fifth convolutional blocks were 64x64, 128x128, 256x256, 512x512, and 512x512, respectively, yielding 610,304 values in total for each visual stimulus.

The bottom row of Fig. 5 shows style synthesized images sharing only style information with the original image. For each image, white noise was used to compute content loss, and the original image was used to compute style loss. The style-synthesized images reproduce almost exactly the surface properties of natural surfaces, including material and gloss. Therefore, image statistics and style information may play such an important role in the perception of surfaces that they can reconstruct images that accurately reproduce surface materials and properties.

#### Reverse-correlation analysis between VEPs and statistical image features

Demonstrations in Fig.5 suggest that statistical features of images, which are thought to be represented even in the VEPs, strongly support the perception of natural surfaces. To verify the image statistics and style information are actually correlated with the VEPs, we performed a reverse correlation analysis between representational similarity of VEPs and the image statistics, or the style information. The VEPs of 29 electrodes (same as Part 1) within 50 ms time-window were converted to the dissimilarity matrix, and we calculated the correlation coefficient between the dissimilarity matrix of the VEPs within each time window and that of the image statistics or style information.

For the analysis, the VEPs of 29 electrodes, excluding Fp1 and Fp2, were used. To clarify how the correlations changed in accordance with latency, we used data from 1∼50, 26∼75, …, 451∼500 ms after stimulus onset. Correlation coefficients were calculated between the dissimilarity matrix of the VEPs and that of image statistics or style information.

### Results

Fig. 6 shows the temporal development of the correlations (*N* = 18,145) between the dissimilarity matrix of the VEPs and that of the image statistics (yellow line) or style information (green line). The correlation with the image statistics reached a maximum at around 200 ms after the stimulus onset (*r*_max_ = 0.22, *p* < .001), while the correlation with style information reached a maximum at around 175 ms after the stimulus onset (*r* = 0.38, *p* < .001).

**Figure 6.**
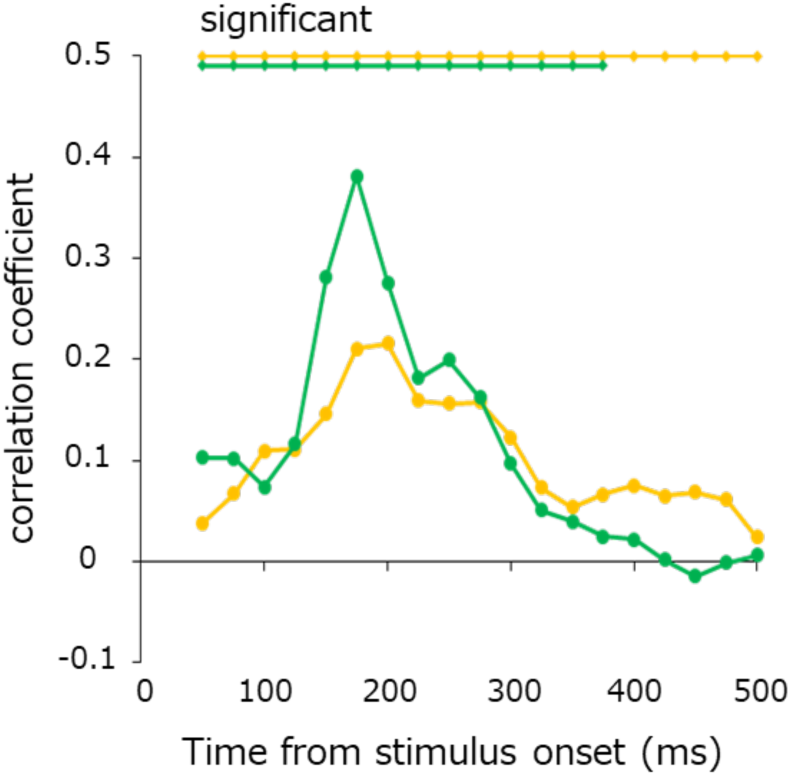
Temporal development of the correlation between the dissimilarity matrix of the VEPs and the image statistics (yellow line), or the style information (green line) for each time window.

Image statistics are spatially global image features computed from the entire image. The style information is also global information because it is the gram matrix, inter-channel correlations, of the activation in the dCNN model (Krizhevsky et al., 2012; Zeiler & Fergus 2014). The dCNN models are thought to consequently learned similar kernels to the receptive field properties of neurons in the human visual cortex, which processes statistical structure of natural images. Therefore, these results suggest that the perception of natural surfaces is strongly supported by statistical image features, which are highly reflected on the VEPs.

## Part 3: Surface Image Reconstruction from VEPs

The results of Part 1 and Part 2 suggest that the perception of natural surfaces is supported by statistical image features, and such features are strongly reflected in the VEPs for natural surface images. Inspired by these results, we hypothesized that it might be possible to reconstruct statistical features from VEPs and even decode perceptual impressions of natural surfaces based on these features. In Part 3, we developed a multimodal variational autoencoder (MVAE) model (Fig. 7) using the VEPs (up to 500 ms after the stimulus onset) measured in Part 1 and their corresponding visual stimuli as inputs, and we reconstructed the visual stimuli themselves only by the VEPs in the testing data.

**Figure 7.**
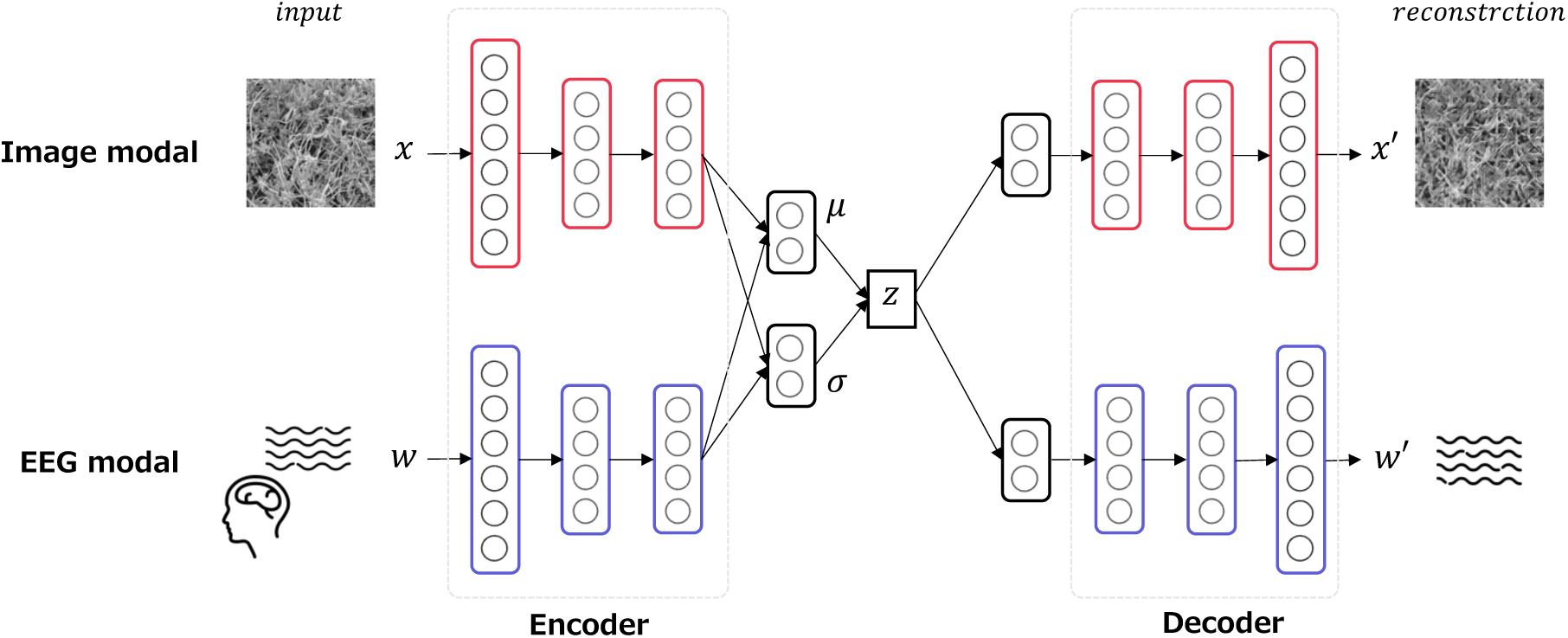
Schematic diagram of the MVAE models.

### Materials & Methods

#### MVAE models

Fig. 7 shows a schematic diagram of the developed MVAE model based on Wakita, Orima, & Motoyoshi (2021). Surface images and VEPs of a certain variant were used as testing data, and the remaining variants were assigned as the training data. 5-fold cross-validations were performed to ensure that each variant was assigned to the testing data (variant condition).

The EEG data were preprocessed as described in the method section of Part 1 and treated as 500 dimensional vector data (from 1 ms to 500 ms after the stimulus onset). 30-35 samples of EEG data corresponding to a single texture stimulus were picked up in random combinations and averaged for input to the model. The average waveforms were normalized to min0 and max1 to eliminate the influence of the absolute value of each channel (cf. Wakita et al., 2021). The VEPs of 29 electrodes (F3, F4, C3, C4, P3, P4, O1, O2, F7, F8, T7, T8, P7, P8, Fz, Cz, Pz, FC1, FC2, CP1, CP2, FC5, FC6, CP5, CP6, FT9, FT10, TP9, and TP10) were used as the EEG modal input. The visual stimuli resized to 128 x 128 pixels was used as the image modal input. The reconstructed image was also a 128 x 128 pixels image. The learning rate was set to 5e-5 and the Adam gradient descent method was used to train the model. The batch size was 16, vector size of latent variables in the MVAE model was 256, and the training epoch was 1,500.

Table 1 shows the architecture of the MVAE model. In practice, except for the final output layer, each convolution layer is followed by batch normalization and ReLU rectifier. The design of the loss function is the same as in the previous study (Wakita et al., 2021).

**Table 1.** Architecture of the MVAE models trained with all VEPs.

| Image modal |  | EEG modal |  |
| --- | --- | --- | --- |
| encoder | decoder | encoder | decoder |
| Input 3x128x128 | FC 256-256 | Input 29x500 | FC 256-256 |
| Conv2d 1-32-1 | UpConv2d 3-512-1 | Conv1d 3-128-2 | UpConv1d 3-256-2 |
| ResNetBlock 3-32-1 | ResNetBlock 3-512-1 | ResNetBlock 3-128-1 | ResNetBlock 3-256-1 |
| AvgPool2d 1/4 | UpConv2d 2-256-1 | Conv1d 1-128-1 | UpConv1d 3-128-2 |
| Conv2d 1-64-1 | ResNetBlock 3-256-1 | ResNetBlock 3-128-1 | ResNetBlock 3-128-1 |
| ResNetBlock 3-64-1 | UpConv2d 2-128-1 | Conv1d 1-128-1 | UpConv1d 3-128-2 |
| AvgPool2d 1/2 | ResNetBlock 3-128-1 | ResNetBlock 3-128-1 | ResNetBlock 3-128-1 |
| Conv2d 1-128-1 | UpConv2d 4-64-1 | Conv1d 1-128-1 | UpConv1d 3-64-2 |
| ResNetBlock 3-128-1 | ResNetBlock 3-64-1 | ResNetBlock 3-128-1 | ResNetBlock 3-64-1 |
| AvgPool2d 1/2 | UpConv2d 4-32-1 | AvgPool1d 1/3 | UpConv1d 3-64-2 |
| Conv2d 1-256-1 | ResNetBlock 3-32-1 | Conv1d 3-256-2 | ResNetBlock 3-64-1 |
| ResNetBlock 3-256-1 | UpConv2d 1-3-1 | ResNetBlock 3-256-1 | UpConv1d 3-32-2 |
| AvgPool2d 1/2 | Sigmoid | Conv1d 1-256-1 | ResNetBlock 3-32-1 |
| COnv2d 1-512-1 | Output 3x128x123 | ResNetBlock 3-256-1 | UpConv1d 3-32-2-2 |
| ResNetBlock 3-512-1 |  | Conv1d 1-256-1 | ResNetBlock 3-32-1 |
| Conv2d 1-512-1 |  | ResNetBlock 3-256-1 | UpConv1d 4-29-2-2 |
| AvgPool2d 1/4 |  | Conv1d 1-256-1 | ResNetBlock 3-29-1 |
| FC 512-256 (mean) |  | ResNetBlock 3-256-1 | UpConv1d 1-29-1 |
| FC 512-256 (var) |  | AvgPool1d 1/3 | ResNetBlock 3-29-1 |
|  |  | Conv1d 3-512-2 | Sigmoid |
|  |  | ResNetBlock 3-512-1 | Output 29x500 |
|  |  | Conv1d 1-512-1 |  |
|  |  | ResNetBlock 3-512-1 |  |
|  |  | Conv1d 1-512-1 |  |
|  |  | ResNetBlock 3-512-1 |  |
|  |  | Conv1d 3-256-2 |  |
|  |  | AvgPool1d 1/2 |  |
|  |  | FC 256-256 (mean) |  |
|  |  | FC 256-256 (var) |  |

**Table 2.**
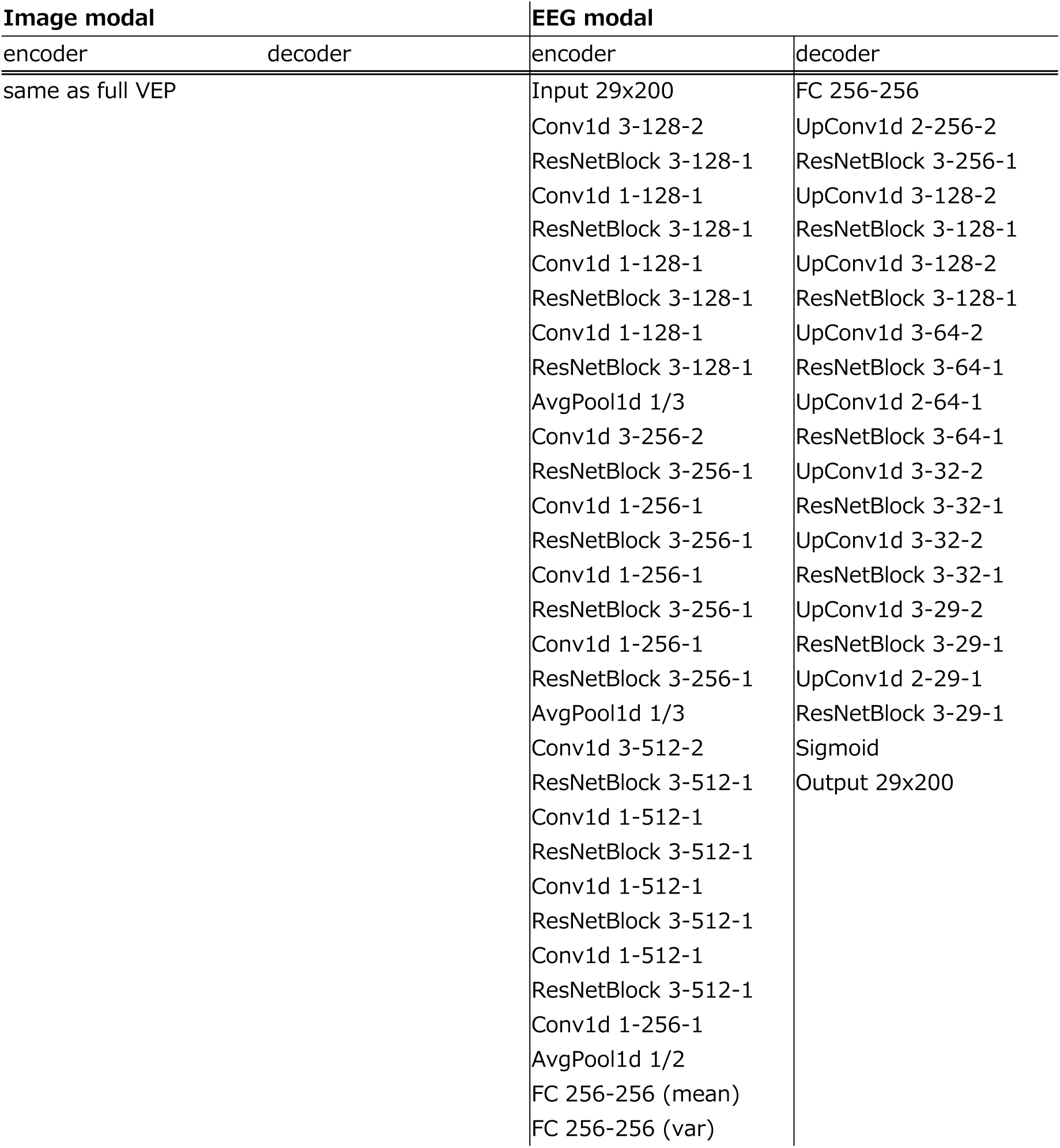
Architecture of the MVAE models trained with VEPs within 200 ms from the stimulus onset.

| **Image modal** |  | **EEG modal** |  |
| --- | --- | --- | --- |
| encoder | decoder | encoder | decoder |
| same as full VEP |  | Input 29x200 | FC 256-256 |
|  |  | Conv1d 3-128-2 | UpConv1d 2-256-2 |
|  |  | ResNetBlock 3-128-1 | ResNetBlock 3-256-1 |
|  |  | Conv1d 1-128-1 | UpConv1d 3-128-2 |
|  |  | ResNetBlock 3-128-1 | ResNetBlock 3-128-1 |
|  |  | Conv1d 1-128-1 | UpConv1d 3-128-2 |
|  |  | ResNetBlock 3-128-1 | ResNetBlock 3-128-1 |
|  |  | Conv1d 1-128-1 | UpConv1d 3-64-2 |
|  |  | ResNetBlock 3-128-1 | ResNetBlock 3-64-1 |
|  |  | AvgPool1d 1/3 | UpConv1d 2-64-1 |
|  |  | Conv1d 3-256-2 | ResNetBlock 3-64-1 |
|  |  | ResNetBlock 3-256-1 | UpConv1d 3-32-2 |
|  |  | Conv1d 1-256-1 | ResNetBlock 3-32-1 |
|  |  | ResNetBlock 3-256-1 | UpConv1d 3-32-2 |
|  |  | Conv1d 1-256-1 | ResNetBlock 3-32-1 |
|  |  | ResNetBlock 3-256-1 | UpConv1d 3-29-2 |
|  |  | Conv1d 1-256-1 | ResNetBlock 3-29-1 |
|  |  | ResNetBlock 3-256-1 | UpConv1d 2-29-1 |
|  |  | AvgPool1d 1/3 | ResNetBlock 3-29-1 |
|  |  | Conv1d 3-512-2 | Sigmoid |
|  |  | ResNetBlock 3-512-1 | Output 29x200 |
|  |  | Conv1d 1-512-1 |  |
|  |  | ResNetBlock 3-512-1 |  |
|  |  | Conv1d 1-512-1 |  |
|  |  | ResNetBlock 3-512-1 |  |
|  |  | Conv1d 1-512-1 |  |
|  |  | ResNetBlock 3-512-1 |  |
|  |  | Conv1d 1-256-1 |  |
|  |  | AvgPool1d 1/2 |  |
|  |  | FC 256-256 (mean) |  |
|  |  | FC 256-256 (var) |  |

For comparison, another type of MVAE models were developed using only the VEPs within 200 ms from the stimulus onset. This was because it would be interesting to compare this model with the MVAE model trained using the full length VEPs as input, since the results of Part 1 suggested that the encoding of information related to low-level image features and surface properties is completed within 200 ms from the stimulus onset. This MVAE model was trained under the same conditions as the MVAE model developed first, but because the size of the input VEPs was different, the structure of the model was slightly different. We developed both MVAE models using almost the same number of parameters as possible.

### Results

#### Reconstruction of visual stimuli from the VEPs (variant condition)

Under the variant condition, train/test split was performed in accordance with the five types of variants of surface images cropped from each original image. Four variants were assigned to the training set, and remaining one variant was assigned to the testing set. The MVAE model was trained by the images and VEPs in the training set, and the reconstructed images were generated only by the VEPs in the testing set. We performed another type of train/test split (image-split condition), which we will discuss later.

Fig. 8a shows the reconstructed images when the VEPs in the testing data were input to the MVAE model, which has been trained with the full length VEPs in the training set. The input VEPs were normalized to min0 and max1 for each image and within each channel in the same way as training, and then averaged. As shown in Fig. 8a, reconstructed images are similar to the original images, and some of them were photorealistic and hard to distinguish from the original images.

**Figure 8.**
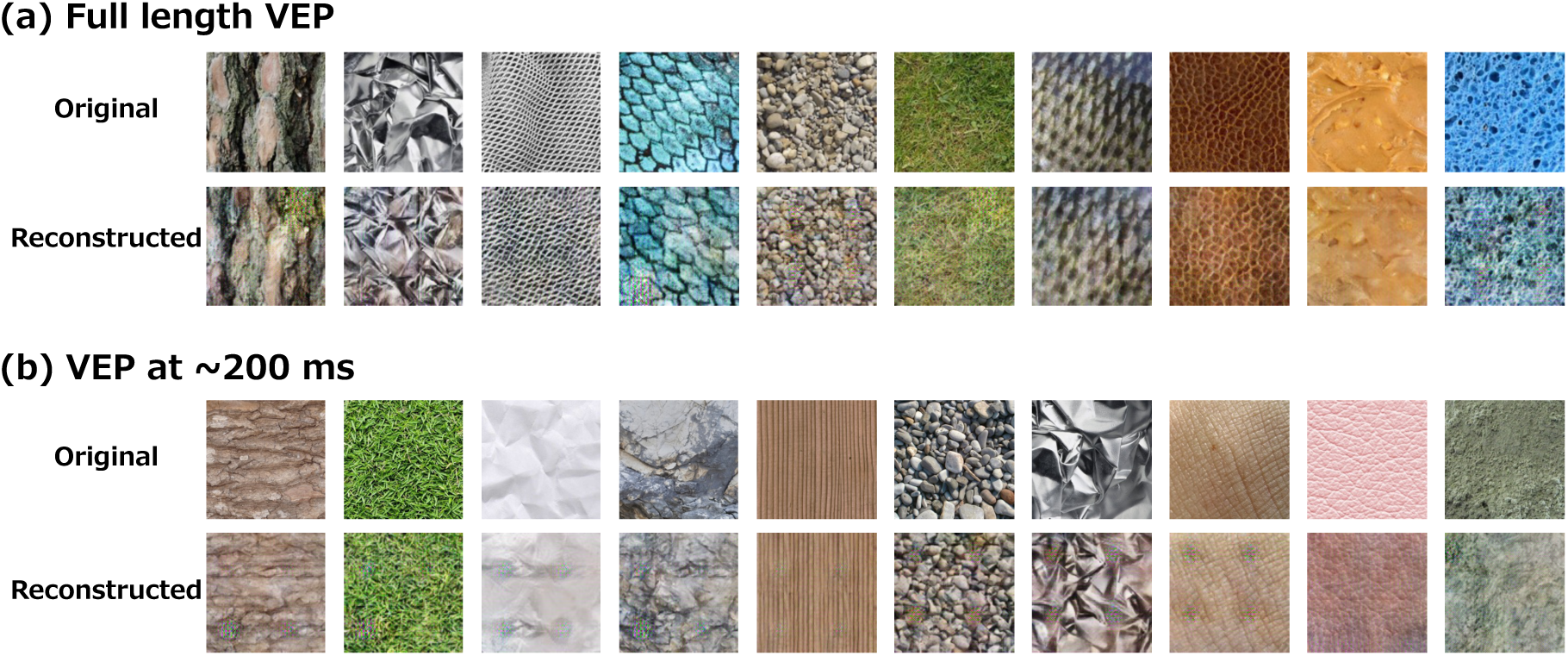
Examples reconstructed images only from the VEPs in the testing data (variant condition). (a) Reconstruction images via the MVAE models trained with full length VEPs, (b) Reconstruction images via the MVAE model trained with VEPs within 200 ms from stimulus onset.

Fig. 8b shows the reconstructed images when the VEPs in the testing data were input to the MVAE model, which has been trained with the VEPs within 200 ms from the stimulus onset. Although some reconstructed images were comparable to those in Fig. 8a, many of them had lower quality.

To quantitatively assess the quality of the reconstructed images, we examined whether the reconstructed images in each validation set were similar to the original images through psychophysical rating experiments. We adopted the two alternative forced choice (2AFC) experiment, following the previous study (Wakita et al., 2021). In the experiment, the original image was presented at the center of the screen, the corresponding reconstructed image (target) was presented on one side, and a randomly selected non-corresponding reconstructed image (non-target) was presented on the other side. Eleven observers responded which image was more similar to the original image by pressing a button with free viewing. The viewing distance was adjusted so that the stimulus size was 5.7 x 5.7 deg (40 cm). Observers were strongly instructed to judge by pure visual appearance, not by categorical classification. Responses were made once for each validation set, and the responses to the five validation sets were averaged to obtain a representative value for each observer. Finally, the percentage of correct responses was averaged across observers. The reconstructed images from the two types of MVAE models were evaluated in separate experimental blocks. Eleven observers participated in the experiment because of the experimental schedule.

Fig. 9a shows the results of the experiment. The dots at the top of each panel indicates that the correct response rate was statistically significant (two-tailed t-test; *p* <= .031, |*t*(10)| >= 2.52 for 500 ms, *p* <= .027, |*t*(10)| >= 2.59 for 200 ms). We adopted the BH FDR correction method with α of .05 to address the multiple comparison.

**Figure 9.**
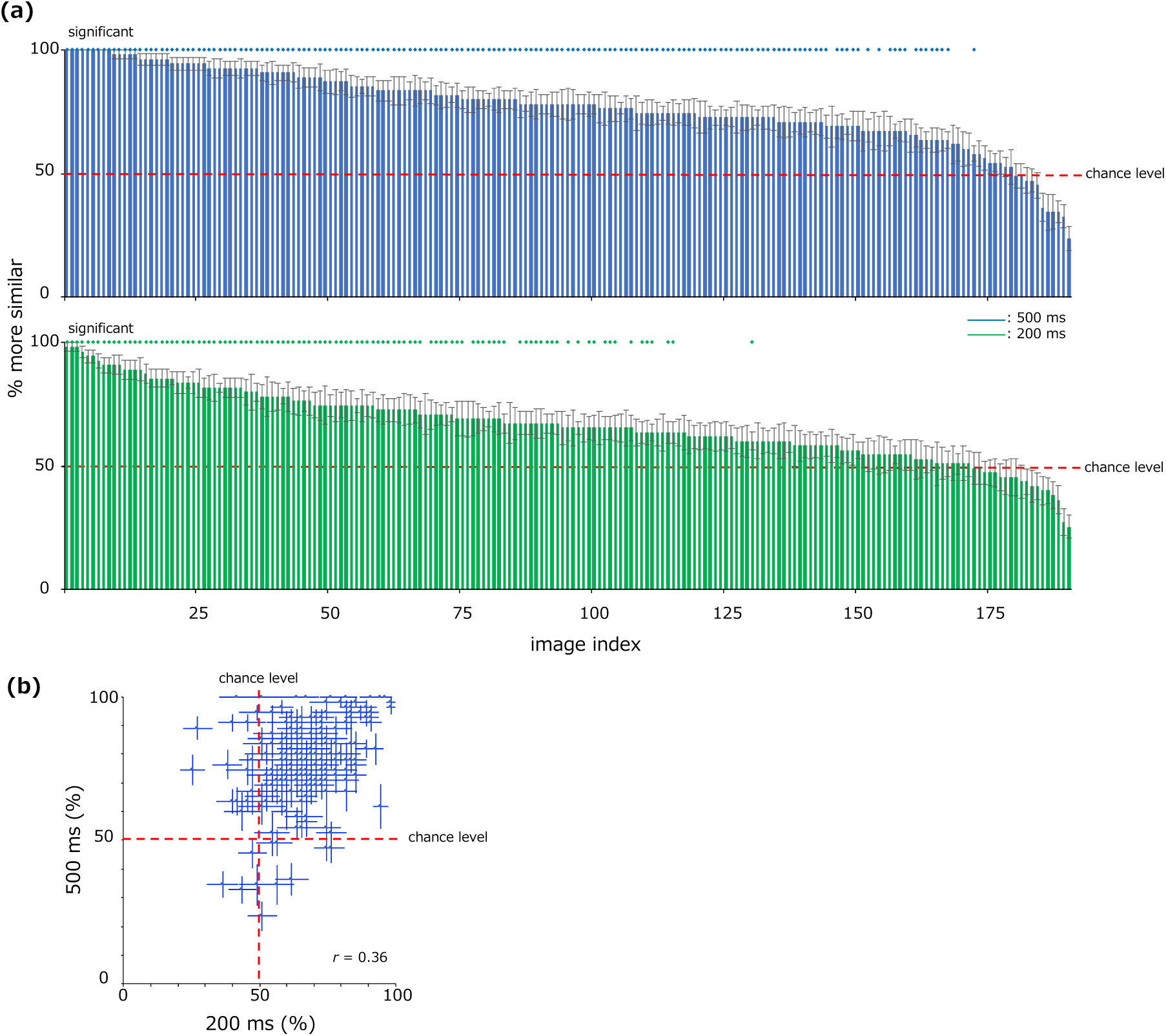
(a) Probability of responses that the reconstructed images (target) were more similar to the original images than a non-target image. Blue indicates the results for the full length VEPs, and green indicates the results for the VEPs within 200 ms from the stimulus onset. The red dotted line indicates chance level, and the dots at the top of each panel indicates that the correct response rate was statistically significant. (b) A scatter plot of the data in (a). The horizontal axis represents the reconstruction from the VEPs with 200 ms from the stimulus onset, and the vertical axis represents the reconstruction from the full length VEPs. The red dotted lines indicate chance levels, and error bars at each point indicate ±1 s.e.m. between observers in the corresponding axes.

We found that 54.5 % of reconstructed images from the VEPs within 200 ms from the stimulus onset were more similar to the original images (green bars), while 85.9 % of the reconstructed images from the full length VEPs were more similar to the original images (blue bars). We note that we considered a target image is ‘more similar’ than a non-target image if the correct response rate was significantly above the chance level (50 %) of the 2AFC experiment.

These results suggest that the perceptual appearance of the surface image itself was successfully reconstructed from the VEPs. As shown in the scatterplot in Fig. 9b, the correlation between the ratings was 0.36. Overall, the reconstructed images from the VEPs within 200 ms from the stimulus onset had lower quality, but we could not find consistent trend across conditions.

In addition, for the MAVE model trained on the full length VEPs, we also tested which material category the reconstructed images would be classified into. Ten observers classified the reconstructed images presented at the center of the screen into one of 20 categories according to the legend presented on the right side of the screen. Responses were made once for each validation set, and the proportion of correct responses to the five validation sets were averaged to obtain a representative value for each observer. Finally, the proportion of correct responses was averaged across observers. As shown in Fig. 10a, for 69.6 % of the images, the reconstructed images were classified into the same material category as the original images at a statistically significant level (two-tailed t-test; *p* <= .027, |*t*(9)| >= 2.65), suggesting that the material category was successfully reproduced by the VEPs. Fig. 10b shows the average proportion of correct for each material category. The results show that the images of grass, fish scale, paper, gravel, and leather were particularly well reconstructed, and all material categories were reconstructed at statistically significant level (*p* <= .022, |*t*(9)| >= 2.74). We note that we adopted the BH FDR correction method with α of .05 to address the multiple comparison.

**Figure 10.**
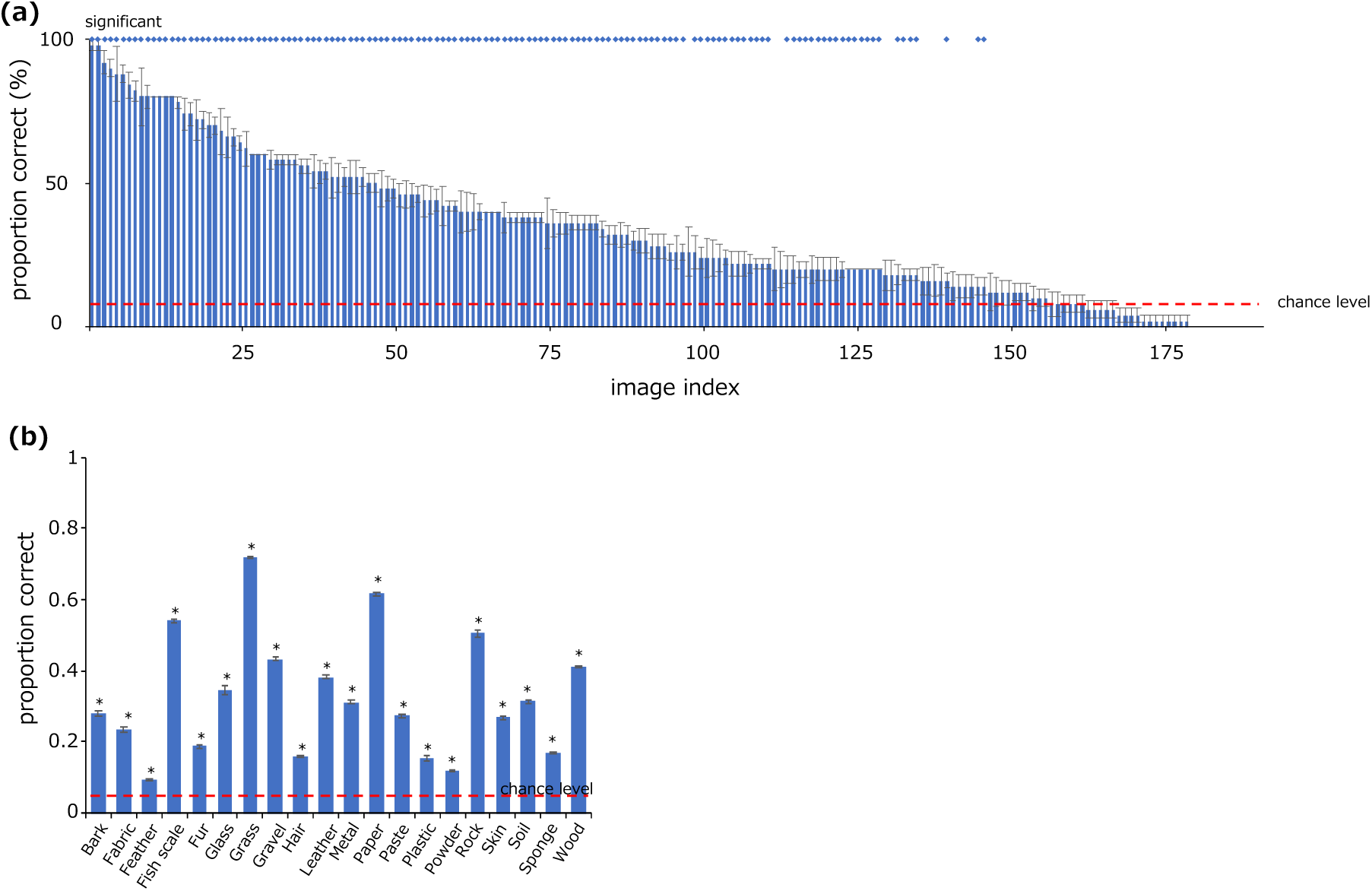
(a) Probability of responses that the reconstructed images were classified into the correct material category. The experiment was conducted only for the reconstructed images from the full length VEPs. (b) Average probability of responses for each material category that the reconstructed images were classified into the correct material category. In each panel, the red dotted line represents the chance level, the dots in (a) and the * in (b) at the top of each panel indicate that the proportion of correct was statistically significant. Error bars at each point indicate ±1 s.e.m. between observers.

Furthermore, to quantify the reconstruction quality of perceived surface properties, we conducted the psychophysical rating experiment for the reconstructed images. Nine observers rated the 6 surface properties for the reconstructed images in the same way as in Part 1. Fig. 11 shows the comparison of the surface property ratings for the reconstructed image from the VEPs with those for the original images. The scatter plot in Fig. 11a shows the relationship between the surface property ratings for the original images (X-axis) and that for the reconstructed images (Y-axis) under the variant condition. Together with the results in Fig. 10a, this plot shows that observers perceived similar surface properties in the reconstructed image as in the original image, but the degree of similarity was different for each surface property. Fig. 11b shows the correlations between surface property ratings for the original and reconstructed images: *r* = 0.76, 0.66, 0.76, 0.59, 0.62, and 0.61 (*p* < .001) for lightness, colorfulness, smoothness, glossiness, hardness, and heaviness, respectively. The correlations were relatively low for glossiness, hardness, and heaviness.

**Figure 11.**
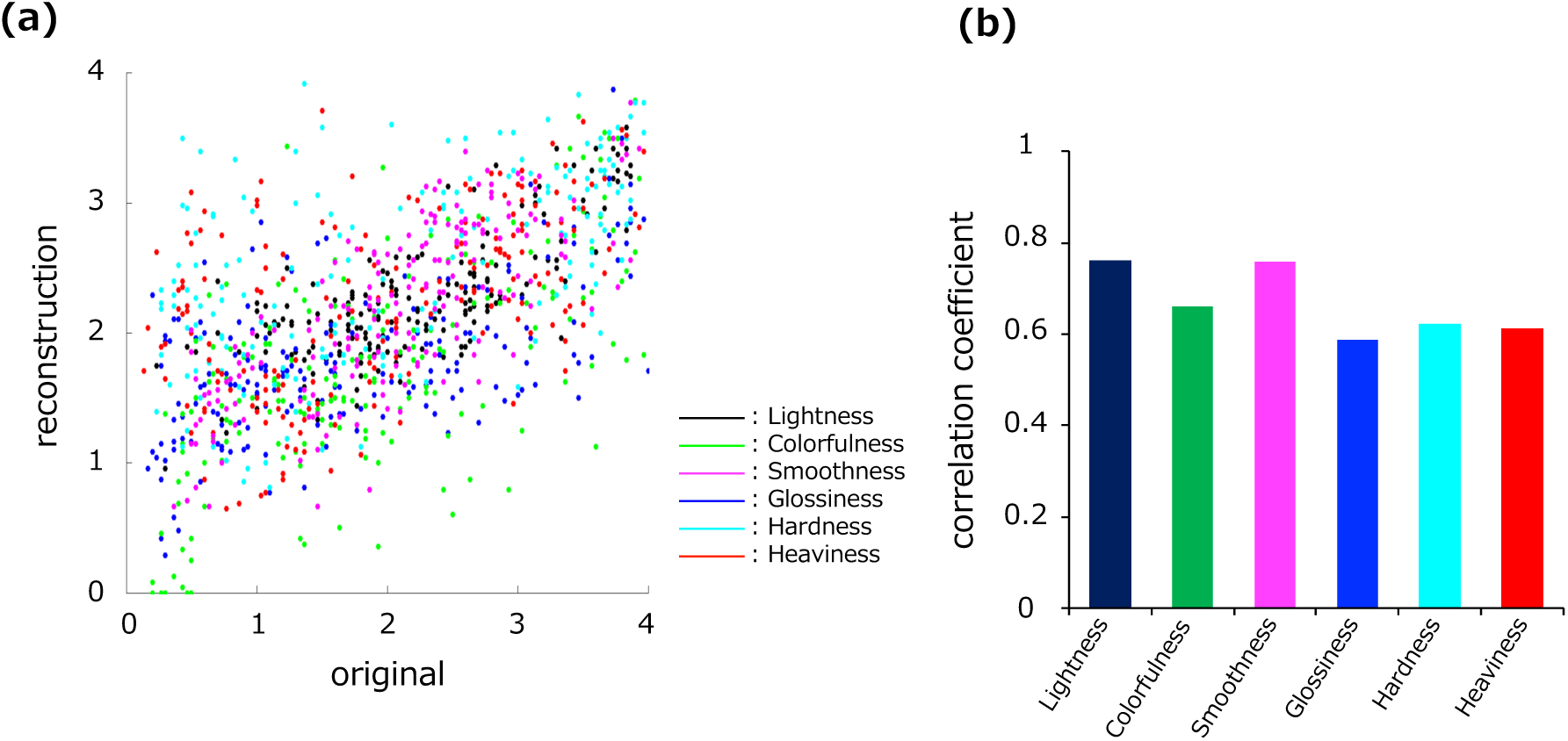
(a) A scatter plot of surface property ratings for the original images and the reconstructed images. Each point corresponds to each image, and each color corresponds to each surface property shown in the right of the panel. (b) Correlation coefficients between the surface property ratings for the original images and the reconstructed images.

We note that the grading of material categories and surface properties was not performed for the MVAE model trained with the VEPs within 200 ms from the stimulus onset, where the reconstruction of the visual stimuli was not successful very much.

#### Classification of material categories and surface properties by latent variables in MVAE models

The reconstructed images from the VEPs retained the material categories and surface properties of the original images. This may be because the MVAE models trained by the visual stimuli and the corresponding VEPs acquired information related to the material categories and surface properties that were reflected in the VEPs. To confirm this hypothesis directly, we tested whether the 256-dimensional latent variables (z) of the MVAE models that was obtained by inputting the VEPs in the testing data could classify the material categories and surface properties of the corresponding visual stimuli. Latent variables were extracted from each of the five trained MVAE models in the variant condition, and each of them was used as the input to SVM models to classify material categories and surface properties of corresponding visual stimulus. The training and testing procedures of the SVM models were the same as Part 1.

The classification accuracy was averaged across the five MVAE models, and finally, a binomial test was conducted using the number of images as the sample size.

Fig. 12 shows the classification accuracy of material categories and surface properties (lightness, colorfulness, smoothness, glossiness, hardness, and heaviness) by the latent variables of the MVAE model. The material categories and all surface properties were significantly classified (*p* < .001, *t*(190) = 10.27, 7.51, 6.41, 7.95, 3.95, 6.96, and 6.58, respectively), and especially smoothness was classified with high accuracy. The result was consistent with our observation that the smoothness of the original images was well reflected in the reconstructed images.

**Figure 12.**
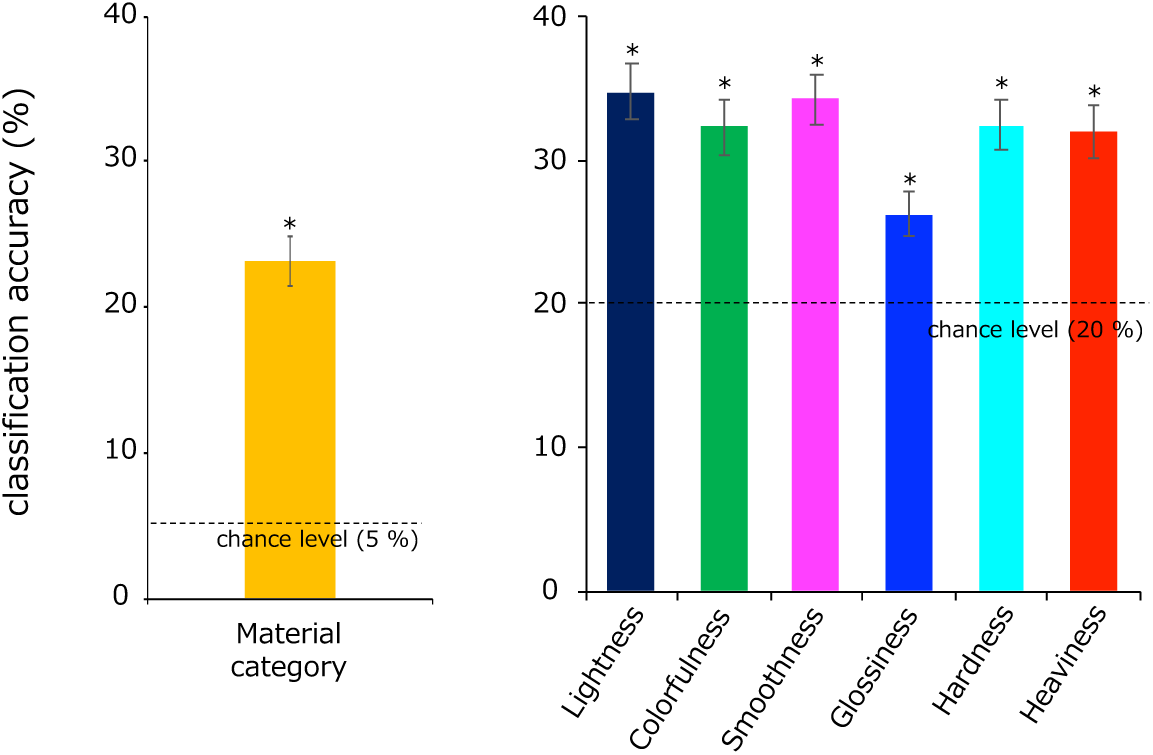
Classification accuracy of material categories and surface properties by the latent variables in the MVAE model. Dotted lines in each panel represent the chance level of the classifications. Error bars indicate ±1 s.e.m. between the 191 images and * indicate the significant classification accuracy.

#### Reconstruction of visual stimuli from VEPs (image-split condition)

The results of variant condition were obtained using only the VEPs in the testing set, but there was a problem that the corresponding visual stimulus of the VEPs in the testing set and those of the VEPs in the training set share the same original images. In this case, under the variant condition, we could not confirm whether the MVAE model can reconstruct images only by the VEPs for perfectly ‘unknown’ images.

To address this problem, we constructed another MVAE model under the condition that the images and corresponding VEPs used for training were not used for testing at all. In this model, 191 original images were split into a 5:1 ratio, and the smaller portion of images were allocated as the testing data (image-split condition). The images included in the testing data and the VEPs for the variant images in the testing data were not used for training at all.

Fig. 13 shows the reconstructed images of the image-split condition. In the image-split condition, the visual stimuli were reconstructed realistically.

**Figure 13.**
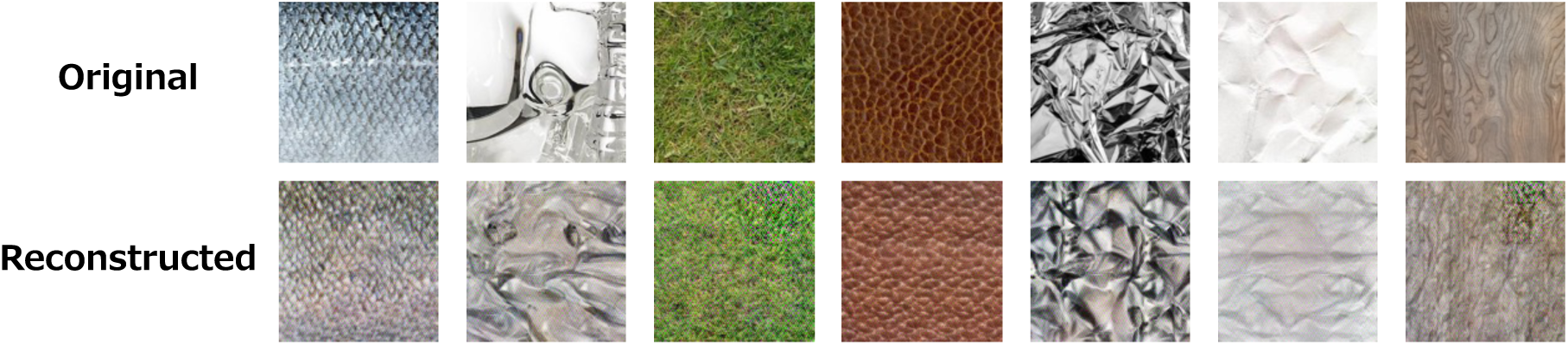
Reconstructed images only from the VEP in the testing data under the image-split condition.

To evaluate the quality of these reconstructed images and to assign material categories, we conducted the rating experiment using the same experimental paradigm as in the variant condition. Ten observers participated in the experiment. As shown in Fig. 14a, 50 % of the reconstructed images were considered more similar to the original images at a statistically significant level (two-tailed t-test; *p* <= .003, |*t*(9)| >= 4.00). In addition, Fig. 14b shows that 28.1% of the reconstructed images were classified into the same material category as the original image at a statistically significant level (two-tailed t-test; *p* <= .024, |*t*(9)| >= 2.70). We note that images with a 0 % correct response rate in Fig. 14b were classified into a specific wrong category, resulting in 0 % accuracy instead of the chance level of 5 % and that we adopted the BH FDR correction method with α of .05 to address the multiple comparison.

**Figure 14.**
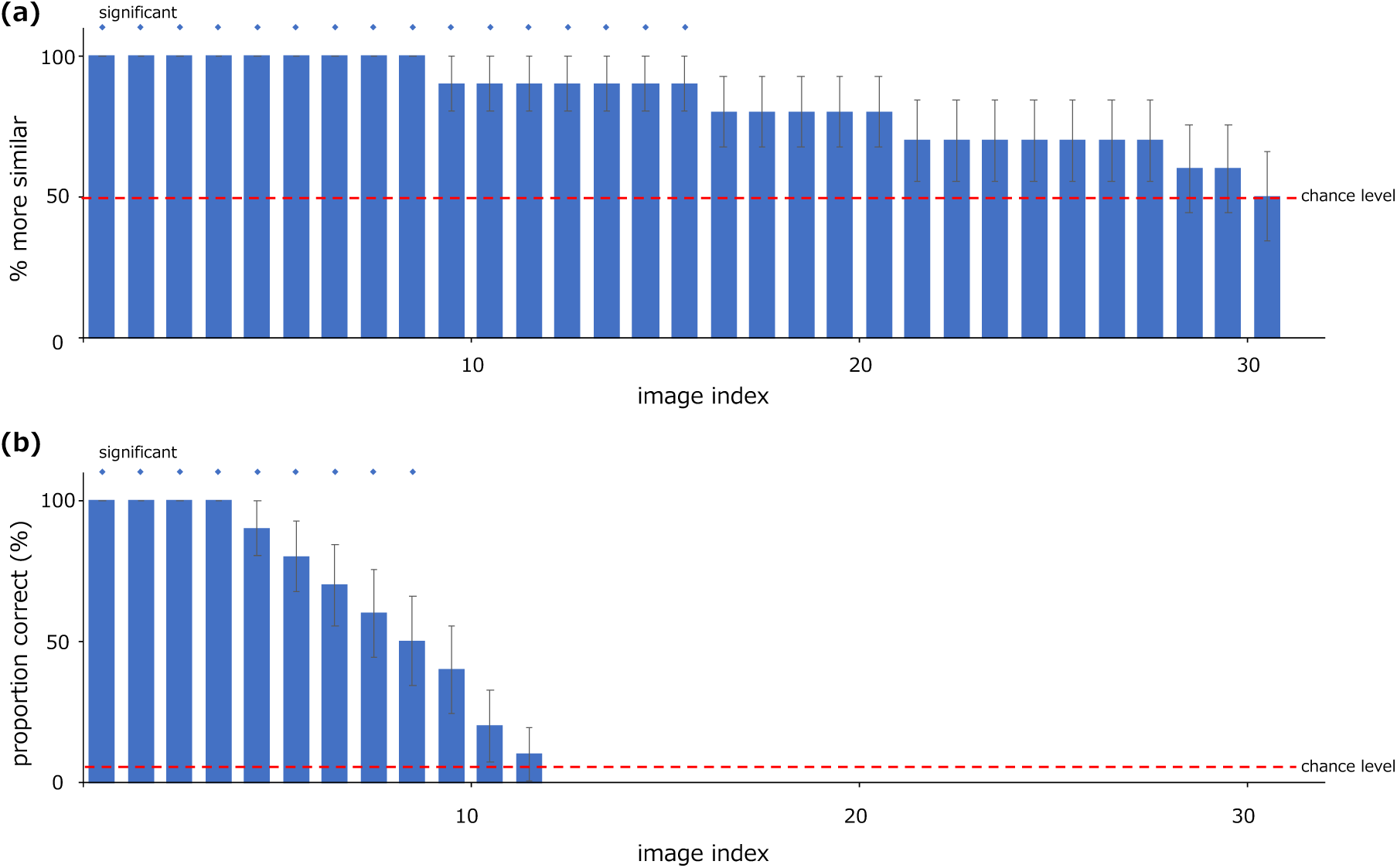
Probability of responses that the reconstructed images (target) were more similar to the original images than a non-target image under the image-split condition. (a) Similarity to the original image, (b) Material category assignment, respectively. In each panel, the red dotted line indicates the chance level, and the blue dots at the top of each panel indicates that the proportion of correct responses was statistically significant. Error bars at each point indicate ±1 s.e.m. between observers.

The results of Part 3 showed that it is possible to reconstruct the surface image itself from the corresponding VEPs for natural surface images, and some reconstructed images were photorealistic. Furthermore, the results of the image-split condition showed that even completely unknown surface images can be reconstructed by the VEPs.

These results in Part 3 suggest that the perception of object surface material and surface properties are strongly based on global textural features to the extent that the visual stimuli themselves can be reconstructed from VEPs for surface images.

### Discussion

In the present study, we measured VEPs for natural surface images made of a wide range of materials to investigate the dynamic neural representations underlying the classification of material categories, surface properties, and the perceptual appearance of natural surfaces. Part 1 showed that material categories and various perceived surface properties (lightness, colorfulness, smoothness, glossiness, hardness, and heaviness) can be decoded from VEP signals with high accuracy using SVM. Part 2 showed that these VEP signals at early latencies strongly correlated with dCNN style information in each image, suggesting that cortical representations of low- and high-level statistical image features play a significant role in the perception of material categories and surface properties. Inspired by these results, in Part 3, we attempted to reconstruct the surface image itself by training deep generative models (MVAE models) using the surface images and corresponding VEPs as input. As a result, some of the reconstructed images remarkably resembled the original images, which demonstrate the decoding of indescribable impression of natural surface images from VEPs. These results further support the idea that the perceptual quality of complex natural surfaces is supported by neural representations of statistical image features that are reflected even in VEPs with low spatial resolution.

#### Rapid neural coding of surface materials

The SVMs trained to classify the material categories and surface properties of the corresponding visual stimuli by the VEPs achieved significant classifications for all conditions. The 20 material categories began to be significantly classified at around 150 ms after stimulus onset, and the classification accuracy reached a maximum at around 175-225 ms under the time-window condition. These results are consistent with previous studies showing that object surface material can be classified from early latency ERPs (Wiebel et al., 2014), some of natural surface material categories are distinguished and perceived based on statistical features of images that are suggested to be encoded in early latency (Orima & Motoyoshi, 2021), and that material categories can be classified by visual stimuli presented for very short periods of time (Sharan et al., 2014).

Another type of SVM significantly classified 6 types of perceived surface properties of each visual stimulus. Lightness and smoothness were classified within 150 ms from the stimulus onset, and colorfulness was classified within 175 ms from the stimulus onset. Especially, smoothness was classified with 70 % of classification accuracy, which was much higher than the chance level. These results suggest that lightness, colorfulness, and smoothness can be highly characterized only by the statistical image features that are easily reflected in the VEP at early latency (Orima & Motoyoshi, 2021). In fact, lightness, colorfulness, and smoothness correlates very strongly with the pixel mean of the luminance channel, the mean or SD of the color channel, and the meddle to high spatial frequency subband SD, respectively. These surface properties are also controlled by the manipulation of these image statistics (Motoyoshi et al., 2007; Giesel & Zaidi, 2013), suggesting that it is highly likely that these surface properties can be described by image statistics.

On the other hand, although the perceived glossiness, hardness, and heaviness of the surface images could be classified at a statistically significant level, the classification accuracy by the VEPs at early latencies remained lower than the other surface properties and became significant continuously for the first time at 200 ms after the stimulus onset. These results suggest that glossiness, hardness, and heaviness were perceived based on information whose encoding process is not easily reflected in the VEPs at early latencies. Previous studies have suggested that surface glossiness can be manipulated by the skewness of subband images (Motoyoshi et al., 2007), but it seems that its encoding was not very early in the visual cortex. In fact, skewness can be computed by combining the outputs of simple and complex cells of V1, but it is difficult to describe it only by their respective outputs. Therefore, the fact that only late latency VEPs for extremely glossy surface images could significantly classify the glossiness is not inconsistent with the previous findings. On the other hand, the fact that the glossiness of visual stimuli could be classified from VEPs is consistent with the idea that glossiness can be captured by image features, as suggested by the previous studies that pointed out that the glossiness of visual stimuli can be easily captured (Nagai et al., 2015) and described by image statistics (Motoyoshi et al., 2007). In the present study, only extremely glossy stimuli were significantly classified, but it is possible that the glossiness may be classified more accurately if the other image features had been controlled.

It was difficult to classify hardness and heaviness by the VEP at early latency. This result seems natural because hardness and heaviness are defined as mechanical, but not optical, properties of a surface. The previous studies also showed that the perceived surface hardness was not significantly classified from low-level image statistics alone, nor from BOLD signals in the early visual cortex such as V1 and V2 (Jacobs et al., 2014; Baumgartner & Gegenfurtner, 2016). These previous findings suggest that perceived surface hardness is difficult to explain solely by image statistics, which are likely to be reflected in early latency VEPs, and that perceived hardness is not largely reflected in brain activity of the early visual system. On the other hand, it has also been suggested that surface hardness can be estimated merely from visual information (Baumgartner et al., 2013), which is consistent with our findings that hardness could be classified by the VEPs at a statistically significant level.

As shown in the demonstration in Fig. 5, synthetic images that shared only the statistical features of the visual stimulus with the original image, such as PS-synthesized and style-synthesized images, retained surface properties similar to those of the original image. Based on these observations, we confirmed that the statistical features of the visual stimulus are very important for the perception of natural surfaces. Furthermore, reverse correlation analysis between these image features and the VEPs revealed that the VEPs at 175-200 ms after the stimulus onset were strongly correlated with the style information. These latencies overlap with the latency at which the classification accuracy of the material category by the VEPs was maximized in Part 1, suggesting that the style information, which is strongly reflected in the VEPs, contributed significantly to the perception of material categories and surface properties. In addition, the VEPs at 175-200 ms after the stimulus onset were revealed to be correlated with the energy of spatial frequency subbands (Hansen et al., 2011), subband image statistics (Orima & Motoyoshi, 2021), and various types of statistical features (Greene & Hansen, 2020). Therefore, it is implied that such style information is a mid-level feature that might be encoded following lower-level statistical features such as image statistics.

Style information is a feature that can accurately represent many types of natural surface properties. Since the style information of surface images correlated strongly with the VEPs at a particular latency, it may be a relatively global image feature that reflects the statistical regularity of natural images, which may also be reflected in the VEPs. Therefore, even glossiness, which has been considered difficult to predict with low-level statistical features in previous studies (Anderson & Kim, 2009) but successfully captured by the style information (Zhang, Oshima, & Motoyoshi, in preparation), was found to be describable with a higher-level statistical feature. These results support the previous studies (Nishida & Shinya, 1998; Motoyoshi et al., 2007; Sawayama & Nishida, 2018) that have shown that the materials and surface properties of natural surfaces depends on statistical features.

#### Reconstruction of surface appearance from EEG

In Part 3, we successfully reconstructed the visual stimulus only by the simple VEPs for the natural surface images. As shown in Fig. 8, some of the reconstructed images reproduced the impression of the original images quite faithfully, and other images were revealed to be reconstructed similarly to the original images (Fig. 9a). In addition, the observers classified the reconstructed images into the 20 material categories used in the experiment, and the results showed that most of the reconstructed images were classified into the same material categories as the original images at a statistically significant level (Fig. 10b). These results indicate that the material of the visual stimuli was also reconstructed.

The MVAE models used for our reconstruction was trained by inputting a set of visual stimuli and their corresponding VEPs, and we defined the reconstructed image as the output image only from the VEPs in the testing set. However, the train/test split for the image modal in this condition was performed with five different variants that were partially cropped from the original images, that is, although the corresponding visual stimuli to the VEPs in the testing set were not used for training, other parts of the original images that shared certain parts were used for training. In this case, the MVAE models in the variant condition reconstructed the visual stimuli partially based on the distribution of inputting images (Naselaris et al., 2009; Wakita et al., 2021; Takagi & Nishimoto, 2023).

Therefore, from a strict point of view, the train/test split was not perfect in the variant condition because the MVAE models ‘knew’ the images that were similar to the corresponding visual stimulus to the VEPs in the testing set. We therefore conducted a similar analysis (image-split condition) in which both the images and VEPs were completely split into training and testing data, that is, the corresponding visual stimulus to the VEPs in the testing set never be used for training.

The results showed that, as in the variant condition, some reconstructed images were very similar to the original images (Fig. 13), and the 50 % of reconstructed images were similar to the original images at statistically significant levels and 28.1 % of them were classified into the same material categories as the original images (Fig. 14). These results in Part 3 suggest that it is possible to reconstruct perceptually very similar images from the VEPs with low spatial resolution, i.e., to decode the visual impression itself, which is difficult to express in simple terms. The results of the image-split condition also showed that the MVAE models had certain generalization performance.

Under the variant condition, most of the reconstructed images were classified into the same material category as the original images and were shown to be more similar to the original images than the non-target images at a statistically significant level. As shown in Fig. 10b, images in all material categories except for feather were reconstructed significantly above the chance level. In addition, as shown in Fig. 11, the surface properties perceived from the original images and those perceived from the reconstructed images were highly correlated, but the correlation was different for each surface characteristic. Specifically, the correlations were high for lightness, colorfulness, and smoothness, but relatively low for glossiness, hardness, and heaviness. The perceived surface properties that were highly correlated are the same as those reflected in the VEPs at early latency, as suggested in Fig. 4. These results indicate that the images reconstructed by the VEPs retain surface properties that are well reflected in the VEPs at early latencies.

#### Comparison to the other brain decoding studies

Previous brain decoding studies have attempted to decode perceptual content or perception itself from brain activity patterns. Typically, it has been suggested that image attributes such as object category, scene category, orientation, and color of visual stimuli can be decoded based on brain activity measured by fMRI (Cox & Savoy, 2003; Haynes & Rees, 2005; Kamitani & Tong, 2005 Brouwer & Heeger, 2009; Parkes et al., 2009; Peelen et al. 2009; Walther et al. 2009; Said et al. 2010). Although these early studies reconstructed low-level characteristics or image features of visual stimuli, which were not visual impression itself, recent studies have proposed techniques to recover the visual stimuli themselves (Shen et al. 2019a, b). However, fMRI has many negative aspects associated with its cost of use, inconvenience of measurement and analysis, and potential invasiveness, which make it unsuitable for daily use as a brain machine interface (BMI). To overcome such disadvantages, brain decoding using EEG has been investigated recently, which is superior to fMRI in terms of portability, potential invasiveness, and cost (Kaneshiro et al., 2015; Palazzo et al. 2020; Wakita et al., 2021). Some of these studies have succeeded in decoding perceived visual stimuli based on brain activity obtained from EEG, following the development of deep generative models such as the generative adversarial network (GAN, Goodfellow et al., 2014). However, the images generated from the brain activity remain blurred images with low spatial resolution (Palazzo et al., 2017, 2020). On the other hand, our previous studies focusing on style loss (Wakita et al., 2021) and the present study have succeeded in reconstructing images, some of which were indistinguishable from the original images and most of which retained the surface properties of the original images. In particular, the fact that the reconstructed images faithfully reproduced the perceived material categories and surface properties provided the evidence on how much information related to the material categories and surface properties is reflected in the VEPs, which were the main theme to address in the present study. This goal would have been difficult to be achieved with the models proposed in the previous study, which reconstructed blurred visual stimuli that were difficult to evaluate material categories and surface properties.

#### Dynamic neural representations for material perception

The results of the present study suggest that low- and high-level statistical features are crucial for the perception of natural surface images. This idea was confirmed by the facts that the classification accuracy of material categories and some surface properties became significant within 175 ms after the stimulus onset as shown in Fig. 3 and Fig. 4, that style information characterizing surface perception strongly correlated with the VEPs within 200 ms after the stimulus onset, and that the visual impression of surface images were successfully reconstructed only by the VEPs via the MVAE models. These results are consistent with the fact that the global information processed in a single glance contributes to the rough perception of scenes and objects as well as natural surfaces (Potter, 1976; Thorpe et al. 2001; Greene & Oliva, 2009; Greene & Hansen, 2020; Jagadeesh & Gardner, 2022; Orima & Motoyoshi, 2023).

On the other hand, as shown in Fig. 3, the classification accuracy of material categories continued to increase after 200 ms from the stimulus onset, and some surface properties, such as glossiness, hardness, and heaviness, were well classified by the VEPs after 200 ms from the stimulus onset. These results indicate that semantic or abstract information that cannot be fully represented by statistical features alone contribute to the natural surface perception. The ERPs at such latencies have been found as the P300 (Sutton et al., 1965), which is said to reflect the perceived uncertainty, and N400 (Kutas & Hillyard, 1980), which is thought to reflect semantic discrepancies. Such ERP components at around 200-400 ms after the stimulus onset are usually observed in experiments involving psychological tasks that require attention to visual stimuli, and it was difficult to measure them directly in the experimental paradigm of the present study. However, it is worth discussing the fact that the VEPs at such latencies was found to be important. Previous studies (Kutas & Federmeier, 2000; Mudrik et al., 2014) have suggested that N400 reflects the use of semantic memory in language comprehension and responses to contextually strange scenes. Therefore, it is possible that the EEG signals at 200-400 ms from the stimulus onset reflected the processing of the meaning of visual stimuli. Although, to some extent, material categories can be distinguished by image features such as color and contrast, semantic analysis of the visual stimuli would also be necessary for the observers to recognize a particular material completely. In addition, the perception of surface properties such as glossiness, hardness, and heaviness would be reaffirmed by haptic perception (Baumgartner et al., 2013) and analysis with attention, even though there are some aspects that can be predicted from image features alone. Therefore, it is possible that the VEPs at later latencies (200 ms or later) also reflected an encoding process of information that contributed to material recognition, which was different from the information reflected in the VEPs at earlier latencies. Similar findings were also reported in the previous study (Orima & Motoyoshi, 2021), in which a component correlated with ‘unnaturalness’ of synthetic textures was found in the VEPs after 200 ms from the stimulus onset.

#### Use of EEGs for understanding visual processing of natural images

The MVAE models developed in the present study reconstructed very high-quality visual stimuli from simple VEPs alone. However, the realistic images such as reconstructed images obtained in the present study were not generated by the VEPs for natural scene images (obtained in Orima & Motoyoshi, 2023). Previous studies have found that the perception of texture, including natural surface images used in the present study, is largely based on low-level statistical features such as image statistics (Bergen & Adelson, 1988; Heeger & Bergen, 1995; Zipser et al. 1996; Portilla & Simoncelli, 2000; Baker & Mareschal, 2001; Motoyoshi et al. 2007; Freeman & Simoncelli, 2011; Freeman et al. 2013; Ziemba et al., 2019; Orima & Motoyoshi, 2021), but natural scenes have many aspects that cannot be explained by low-level features alone (Groen et al., 2013; Epstein & Baker, 2019; Greene & Hansen, 2020). Information reflected in the EEG signals is global because of the limited spatial resolution of the EEG, which is recorded as the summation of a huge number of neurons’ responses (Nunez & Srinivasan, 2006). This may be the reason why the reconstruction of natural scene images was challenging. Therefore, it is reasonable to assume that the remarkable reconstructions of the present study were limited to texture stimuli.

It is also difficult to completely rule out the possibility that the MVAE model developed in the present study is generalizable only to the image set used in the present study. Although the results of the image-split condition confirm that the realistic reconstruction are not the product of complete overfitting to the image dataset of the present study, it was still imperfect. In the image-split condition, the training data included images similar to those assigned to the testing data (e.g., the training data was set to include grass images because the testing data also included grass images). If a VEP for an image that did not resemble any image in the training data, the reconstructed image would not be similar to the original image. One goal of brain decoding research is to fully reproduce perception from brain activity. The present study could play a part, but to achieve this goal completely, it would be necessary to train on a much larger number of images and VEPs than used in the present study, as the AlexNet (Krizhevsky et al., 2012), which is a great object recognition model.

The architecture of the MVAE models used in the present study were almost the same as the models constructed in the previous study (Wakita et al., 2021), except that the number of parameters and layers were increased to address color images. In the MVAE models used in the previous study and in the present study, one of the two modals was based on a two-dimensional image input, and the other was based on a one-dimensional VEP input. Recent study has developed a model called EEGNet, which yields high accuracy in EEG-based classification (Lawhern et al., 2018). In the other previous study, the EEGNet model is used to visualize spatiotemporal cortical mapping for natural scene perception (Orima & Motoyoshi, 2023). By using a two-dimensional network like EEGNet model instead of a one-dimensional convolutional network, it may be possible to reconstruct the original image more realistically and to visualize the contribution of VEPs to the visual impression in a spatiotemporal manner. It would be interesting to explore more valid models as a means of decoding brain activity in the future.

## Data Availability Statement

The raw data supporting the conclusions of this article will be made available by the authors upon reasonable request.

## Author Contribution

Conceptualization: TO, IM

Data Curation: TO

Formal Analysis: TO

Funding Acquisition: TO, IM

Investigation: TO, IM

Methodology: TO, SW, IM

Project Administration: TO, IM

Resources: TO, SW, IM

Software: TO, SW

Supervision: IM

Validation: TO, SW, IM

Visualization: TO, IM

Writing - Original Draft Preparation: TO, IM

Writing - Review & Editing: TO, SW, IM

